# The RNA-binding protein Igf2bp3 is critical for embryonic and germline development in zebrafish

**DOI:** 10.1101/2020.06.23.167163

**Authors:** Yin Ho Vong, Lavanya Sivashanmugam, Andreas Zaucker, Alex Jones, Karuna Sampath

## Abstract

The ability to reproduce is essential in all branches of life. In metazoans, this process is initiated by formation of the germline, a group of cells that are destined to form the future gonads, the tissue that will produce the gametes. The molecular mechanisms underlying germline formation differs between species. In zebrafish, development of the germline is dependent on the specification, migration and proliferation of progenitors called the primordial germ cells (PGCs). PGC specification is dependent on a maternally provided cytoplasmic complex of ribonucleoproteins (RNPs), the germplasm. Here, we show that the conserved RNA-binding protein (RBP), Igf2bp3, has an essential role during early embryonic development and germline development. Loss of Igf2bp3 leads to an expanded yolk syncytial layer (YSL) in early embryos, reduced germline RNA expression, and mis-regulated germline development. Maternal mutants affecting igf2bp3 exhibit abnormal PGCs and adult igf2bp3 mutants show male biased sex ratios. Therefore, Igf2bp3 is required for normal embryonic and germline development.

## Introduction

In eukaryotes, nascent RNA transcripts are bound in complexes of RNA-binding proteins (RBPs) known as heterogeneous nuclear ribonucleoproteins (hnRNPs) for further processing (Gall, 1956). The purpose of these complexes are to direct the fate of its cargo RNAs, such as splicing (Guil *et al*., 2003; Talukdar *et al*., 2011), capping activity (Gamberi *et al*., 1997), polyadenylation (Nazim *et al*., 2016) and export (Nakielny and Dreyfuss, 1996). The composition of the hnRNPs contributes to the fate of cargo molecules. For example, hnRNPs containing the RNA-binding protein Igf2bp1 impart increased stability of bound RNAs, such as *c-myc* (Weidensdorfer *et al*., 2009; Eisenberg and Jucker, 2012; Samuels *et al*., 2020). Mis-regulation of RNAs through abnormal RBP functions often lead to developmental consequences. For instance, knockout of mouse *Igf2bp1* leads to gross defects in development (Hansen *et al*., 2004) and knockout of the RBP Zar1 in zebrafish leads to abnormalities in sex determination, resulting in male-only fish (Miao *et al*., 2017). Several questions remain regarding how individual RNA-binding proteins contribute to development and the mechanism for their actions in specific cell types.

The germline is one of the earliest cell lineages to be specified across many multicellular organisms (Seydoux *et al*., 1996; Knaut *et al*., 2000). The precursor of the germline is the primordial germ cell (PGC). In mammals, the germline forms in the developing embryo, a process which is induced by signals from neighbouring cells (Lawson *et al*., 1999). In other animals, including fruit flies and zebrafish, a substance termed ‘germ-plasm’ is inherited by the egg from the mother, and contains factors that promote the formation of germ-line cells and development of the gonads.

Primordial germ cells receiving germplasm maintain their fate independently of the surrounding soma, whilst proliferating and migrating to the gonadal ridges (Braat *et al*., 1999). Therefore, germplasm components are tightly regulated, and key components, including ribonucleoprotein complexes, are thought to be essential for correct PGC behaviour (Weidinger *et al*., 2003; Hartung, Forbes and Marlow, 2014; Roovers *et al*., 2018). Germplasm RNAs are post-transcriptionally regulated by factors including microRNAs and RNA-binding proteins. RNA-binding proteins have a wide range of roles in processes such as oocyte maturation (Miao *et al*., 2017; Sun *et al*., 2018) and axis formation (Bontems *et al*., 2009; Kumari *et al*., 2013).

In addition to the functions that RNA-binding proteins have when bound to their target RNAs, the mechanism of the interactions is also important.. For example, Ybx1 protein acts as a translational repressor and localisation factor for RNAs during early zebrafish embryogenesis. Ybx1 interacts with a stem-loop element found in the 3’ UTR of its target transcripts such as *sqt* (Gore *et al*., 2005; Gilligan *et al*., 2011; Kumari *et al*., 2013; Zaucker *et al*., 2018). The DAZ family of proteins, which are critical for germline formation (Haston, Tung and Reijo Pera, 2009), interact with target RNAs via a specific sequence motif, and acts as translational activators (Maegawa *et al*., 2002; Collier *et al*., 2005). More recently, post-transcriptional modifications to RNA are proposed to act as another marker for recognition by RNA-binding proteins. The modifications include 6-methyladenosine (m6A), which is recognised by a host of proteins such as Ythdc2 and Ythdf2, which have been proposed to have a role in germline development (Du *et al*., 2016; Bailey *et al*., 2017; Hsu *et al*., 2017; Zhao *et al*., 2017).

Another group of proteins that are now known to recognise m6A modifications is the Insulin-growth factor 2 mRNA binding protein (Igf2bp) family (Huang *et al*., 2018). Igf2bp family members are nucleocytoplasmic proteins containing two RNA-recognition motifs (RRMs) and two K-Homology (KH) didomains (Wächter *et al*., 2013) that are highly conserved. Igf2bp proteins have many diverse functions. In addition to acting as RNA stabilisers (Huang *et al*., 2018; Ren *et al*., 2020), Igf2bp proteins also have roles in localisation, *Xenopus* Igf2bp3 binds to a *vg1* 3’ UTR element to localise *vg1* transcripts to the vegetal cortex of oocytes (Schwartz *et al*., 1992; Elisha *et al*., 1995; Kwon *et al*., 2002). These observations have subsequently shown Igf2bp protein to play a role in development in *Xenopus* (Yaniv *et al*., 2003), mice (Hansen *et al*., 2004) and zebrafish (Ren *et al*., 2020). However, whether Igf2bp proteins have a role in vertebrate germline development remains unexplored, although many lines of evidence point towards this possibility. In zebrafish oocytes, zebrafish Igf2bp3 is enriched in the germplasm-containing structure, the Balbiani body, (Bontems *et al*., 2009) and *Drosophila igf2bp* has been shown to regulate germ-cell maintenance (Toledano *et al*., 2012).

To determine the role of Igf2bp3 in zebrafish development, we generated and analysed zebrafish mutants affecting *igf2bp3*, and find that maternal Igf2bp3 is essential for proper germline development. Loss of zygotic *igf2bp3* is sufficient to induce a male-sex bias in adult zebrafish. Loss of Igf2bp3 leads to a reduction in canonical germline markers, and maternal *igf2bp3* mutants have a mis-regulated germline, with depletion of primordial germ cells and aberrant PGC behaviour. These results highlight a new function for this family of proteins in germline development and identify a key maternal factor that is essential for germline development, maintenance and PGC migration.

## Methods

### Zebrafish husbandry

Adult zebrafish were kept at the ambient temperature in the University of Warwick aquatics facility in compliance to institutional animal care regulations (Westerfield, 2007), and the UK home office animal welfare regulations. Embryos were collected from pair-wise or pooled intercrosses and incubated at 28.5°C in 0.3X Danieau’s solution with methylene blue (17 mM NaCl, 2 mM KCl, 0.12 mM MgSO4, 1.8 mM, Ca(NO_3_)_2_, 1.5 mM HEPES, pH 7.6). Embryos were manually dechorionated with a pair of Dumont Tweezers #5.

### Affinity Purification, Mass spectrometric analysis and Identification of Igf2bp3

NHS (N-hydroxysuccinimide) activated Sepharose 4 fastflow (GE Healthcare) beads coupled with an RNA aptamer alone control or aptamer fusion (zebrafish ndr1 3’UTR sequences fused to a Tobramycin aptamer) was incubated with 10 mg of 20 min post-fertilisation embryonic lysates, washed and eluted with Tobramycin antibiotic (Hartmuth *et al*., 2002; Ward *et al*., 2011). The eluates were run on 5-16% gradient polyacrylramide gel, washed with deionized water and stained with Coomassie blue for 15 minutes. Protein bands were cut into cubes of ∼1mm. Gel pieces were de-stained twice using 50% ethanol (Thermo Fisher Scientific) in 50 mM ammonium bicarbonate (ABC, Fluka) at 22°C for 15 min and dehydrated with 100% ethanol for 5 min with shaking (650 rpm). Dehydrated gel pieces were reduced with 10 mM DTT (Sigma) at 55°C for 30 min and then alkylated with 55 mM iodoacetamide (Sigma) for 20 min in the dark at 22°C. Samples were then washed with 50% ethanol in 50 mM ABC at 22°C for 15 min and then dehydrated with 100% ethanol for 5 min. The gel pieces were hydrated and incubated with 2.5 ng/μl of trypsin (Promega) in 50 mM ammonium bicarbonate (ABC) overnight at 37°C. Peptides were extracted from gel pieces three times with 5% formic acid in 25% acetonitrile (ACN; Thermo FisherScientific) with 5 minute sonications in a water bath. The supernatants were combined in a fresh vial, dried using a vacuum centrifuge at 40°C, resuspended in 55 μl of of 2.5% acetonitrile containing 0.05% trifluoroacetic acid and sonicated for 30 mins. 20µl of the sample was used for liquid chromatography-mass spectrometry analysis as performed in (Hatano *et al*., 2018). Samples were analysed using reversed phase chromatography with two columns, an Acclaim PepMap µ-precolumn cartridge 300 µm i.d. x 5 mm, 5 μm, 100 Å and an Acclaim PepMap RSLC 75 µm i.d. x 50 cm, 2 µm, 100 Å (Thermo Scientific) was used to separate tryptic peptides prior to mass spectrometric analysis. The columns were installed on an Ultimate 3000 RSLCnano system (Dionex) at 40C. Mobile phase buffer A was composed of 0.1% formic acid and mobile phase B was composed of acetonitrile containing 0.1% formic acid. Samples were loaded onto the µ-precolumn equilibrated in 2% aqueous acetonitrile containing 0.1% trifluoroacetic acid for 5 min at 10 µL min-1, after which peptides were eluted onto the analytical column at 250 nL min-1 by increasing the mobile phase B concentration from 6% B to 37% over 100 min, followed by a 3 minute wash at 80% B and a 10 min re-equilibration at 4% B. Eluting peptides were converted to gas-phase ions by means of electrospray ionization and analysed on a Thermo Orbitrap Fusion (Thermo Scientific). Survey scans of peptide precursors from 375 to 1500 m/z were performed at 120K resolution (at 200 m/z) with a 2×105 ion count target. The maximum injection time was set to 150 ms. Tandem MS was performed by isolation at 1.2 Th using the quadrupole, HCD fragmentation with normalized collision energy of 33, and rapid scan MS analysis in the ion trap. The MS2 ion count target was set to 5×103. The maximum injection time was 200 ms. Precursors with charge state 2–6 were selected and sampled for MS2. The dynamic exclusion duration was set to 40 s with a 10 ppm tolerance around the selected precursor and its isotopes. Monoisotopic precursor selection was turned on. The instrument was run in top speed mode. The acquired tandem mass spectra, as Xcalibur (version 2.2) raw files were analyzed using Max-Quant software v1.6.0.16 (Tyanova et al., 2016a,b) against the UniProtK Danio rerio (UP000000437). Trypsin was specified as the digestion enzyme, with up to 2 missed cleavages, and a parent ion mass tolerance of 4.5 ppm for the initial search, with recalibration enabled. Oxidation of methionine was set as a variable modification and carbamidomethyl of cysteine as a fixed modification for all searches. The MS-MS data were further collated using Scaffold software. Fold change in proteins between the control aptamer and zebrafish ndr1 fusion RNA samples were calculated using emPAI in scaffold. The emPAI is a label-free, relative quantitation of the proteins in a mixture based on protein coverage by the peptide matches in a database search result. Igf2bp3 showed > 4-fold enrichment in ndr1-bound eluates.

### Generation and establishment of *igf2bp3* mutants

Cas9 mutants were generated in the Tu background. Target sequences were verified by PCR sequencing and the conserved *igf2bp3* cDNA sequence was used as a target in ChopChop (Labun *et al*., 2016) to produce the target site “GGCTCCCTTCCTCGTAAAAAG” in exon 1. Embryos were injected with 150 pg Cas9 mRNA and 50 pg sgRNA at the 1-cell stage and raised to adulthood as described (Varshney *et al*., 2015). F0s were outcrossed to identify heterozygous F1s, which were outcrossed again to expand the line before intercrossing to retrieve homozygous mutants.

The *igf2bp3* transgenic insertion alleles were obtained from the Burgess laboratory collection, these mutants were outcrossed over two generations and intercrossed to homozygosity.

### Genotyping mutants

DNA was isolated from fin clips by incubating with lysis buffer (10 mM Tris pH 8.3, 50 mM KCl) with proteinase K (200 μg/mL) at 55°C overnight, followed by inactivation at 95°C for 10 minutes and lysates were used directly for PCR. PCR products were visualised on a 2-3% agarose gel.

The *igf2bp3*_*la020659Tg*_ insertion mutant fish were genotyped by the use of three primers in a single PCR reaction, a forward/reverse primer flanking the insertion and a second reverse primer specific to the long terminal repeats in the retroviral construct. This generates a 250 bp WT product and a ∼800 bp product for the insertion allele. The *igf2bp3*_*la010361Tg*_ insertion allele was genotyped by the use of three primers in a single PCR reaction, a forward/reverse primer flanking the insertion and a second forward primer specific to the long terminal repeats in the retroviral construct. This generates a 350 bp WT product and a ∼900 bp product for the insertion allele. The *igf2bp3*_*Δ 7 bp*_ deletion allele contains a continuous 7 bp deletion in exon 1. This mutation generates a BssaI (NEB) restriction site. PCR products digested by BssaI.

### Synthesis of RNAs

Synthetic capped mRNAs were transcribed using the SP6 mMessage mMachine, following the manufacturer’s instructions. Cas9 sgRNA was transcribed using the T7 HiScribe High Yield RNA Synthesis kit, following the manufacturer’s instructions. RNAs were purified with phenol-chloroform extraction. The pSP64-mmGFP5-nos1-3’UTR (Köprunner *et al*., 2001) and pSP64-eGFP-F-nos1-3’UTR (Weidinger *et al*., 2002) constructs were linearised with SacII and NotI restriction enzymes (NEB) respectively, following the manufacturer’s instructions. These were used to label the cytoplasm and cell membranes of the PGCs.

Probes for whole mount *in situ* hybridisation were produced with Promega SP6/T7/T3 polymerases with DIG-labelling mix (Roche) and purified with lithium chloride extraction.

### Whole mount *in situ* hybridisation

Embryos were fixed in 4% paraformaldehyde in PBS (phosphate buffered saline), and processed to detect gene expression as described previously (Lim *et al*., 2012).

### Total RNA extraction and qRT-PCR

Embryos were collected at various stages and lysed in TRIzol, followed by RNA extraction using the Monarch Total RNAMiniprep kit according to the manufacturers’ instructions. First strand cDNA was synthesised using the PCRBIO cDNA synthesis kit. qPCR samples were prepared with the PCRBIO SyGreen Blue Mix Lo-ROX, and analysis performed with the Stratagene MX3005P.

### Protein gel electrophoresis and Western blot

SDS-PAGE gels were prepared using the Bio-Rad protein electrophoresis systems, zebrafish embryos were homogenised in RIPA (50 mM Tris-HCl, 150 mM NaCl, 1% (v/v) NP-40, 0.5% (w/v) sodium deoxycholate, 1 mM EDTA, 0.1% (w/v) SDS) lysis buffer supplemented with protease inhibitor cocktail (Sigma-Aldrich) using a syringe and needle. Supernatants were collected after a brief centrifugation and boiled in 4X loading buffer (200 mM Tris-HCl (pH 6.8), 400 mM DTT, 8% SDS, 0.4% bromophenol blue and 40% glycerol) before loading.

After transfer of proteins, membranes were rinsed with TBSTw (Tris Buffered Saline, 0.1% Tween-20) once and blocked in 5% skimmed-milk powder in TBSTw for 1 hour before incubation with primary antibody overnight. After incubation, membranes were rinsed 4X in TBSTw for 5 minutes and transferred to secondary antibody for 4 hours at room temperature, excess antibody subsequently removed by a further 4 washes in TBSTw for 5 minutes before detection with ECL Western blotting reagent (Bio-Rad) following manufacturer’s instructions. Signal detection was performed using a ChemiDoc MP Imaging system (Bio-Rad) or with CL-XPosure Film (34089, Thermo Fisher Scientific).

### Image Acquisition

For live tracking of PGCs, embryos were mounted in 0.8% LMPA on a heated stage maintained at 28.5°C and imaged with an Andor Revolution Spinning Disk system, based on a Nikon Ni-E PFS inverted microscope equipped with a Yokogawa CSU-X1 spinning disk unit, fitted with a 488 nm laser and captured with an iXon Ultra 888 EMCCD camera. Images were captured using either a Nikon Plan Apochromat 20X/0.75 NA or the Nikon Apochromat 60X/1.49 NA oil immersion objectives. Images were acquired with the Andor iQ3 software.

For time-lapse imaging of migrating PGCs, images were acquired using a 20X objective and z-stacks were generated with 1 μm step-sizes at 1 minute intervals for 60 minutes. Maximum intensity projections were generated ssing ImageJ/Fiji, and the MTrackJ (Meijering, Dzyubachyk and Smal, 2012) plugin was used to derive displacement, speed and straightness of PGCs.

For analysing PGC filopodia dynamics, z-stacks were generated with the 60X objective with 0.5 μm step-sizes at 10 second intervals for 2-10 minutes. Using ImageJ/Fiji, maximum intensity projections were generated and the filopodia numbers per PGC were recorded by counting the filopodia for a single timepoint. Persistence of the filopodia was calculated by counting the number of consecutive frames that a single filopodium was present. The length of a filopodia was calculated as the average length over its observable lifetime. The frequency of filopodia projections was calculated by measuring the direction of the projections relative to the embryonic midline.

To image PGCs *in vivo* during segmentation, the Zeiss LSM 880 scanning confocal microscope was used.

## Results

### Mutations in *igf2bp3* result in biased sex ratios in adult zebrafish

Igf2bp3 was identified from a proteomic screen to identify RNA-binding proteins in early zebrafish embryonic lysates that bind an RNA aptamer. Igf2bp3 was enriched 4-fold compared to controls. Analysis of *igf2bp3* expression by WISH shows that the transcript is ubiquitously expressed and at high levels during early development (Fig Supp 1A-B). Igf2bp3 protein is expressed maternally and zygotically (Fig Supp 1E), precluding the use of morpholinos to uncover the role of *igf2bp3* in development.

To identify zebrafish lines with genomic lesions in *igf2bp3*, we searched a mutant collection generated by the Lin and Burgess laboratories (Varshney *et al*., 2013). We identified several retroviral transgenic insertion alleles that harbour integration of the *Tg (nLacZ-GTvirus)* in intron one of the *igf2bp3* locus: *igf2bp3*_*la010361Tg*_ and *igf2bp3*_*la020659Tg*_, hereafter referred to as *igf2bp3*_*la361Tg*_ and *igf2bp3*_*la659Tg*_ (Fig 1B). The *igf2bp3* insertion mutant alleles are predicted to result in loss-of-function by preventing splicing of exons one and two in *igf2bp3* transcripts, leading to premature termination codons. Western blots performed using an antibody directed against a central region of Igf2bp3 downstream of the insertion site did not reveal any detectable Igf2bp3 in mutant embryo lysates from both alleles (Fig Supp 1C-E). RT-PCRs of exon-spanning junctions of *igf2bp3* showed a reduction of RNA levels (Fig Supp 1C).

**Figure 1.**
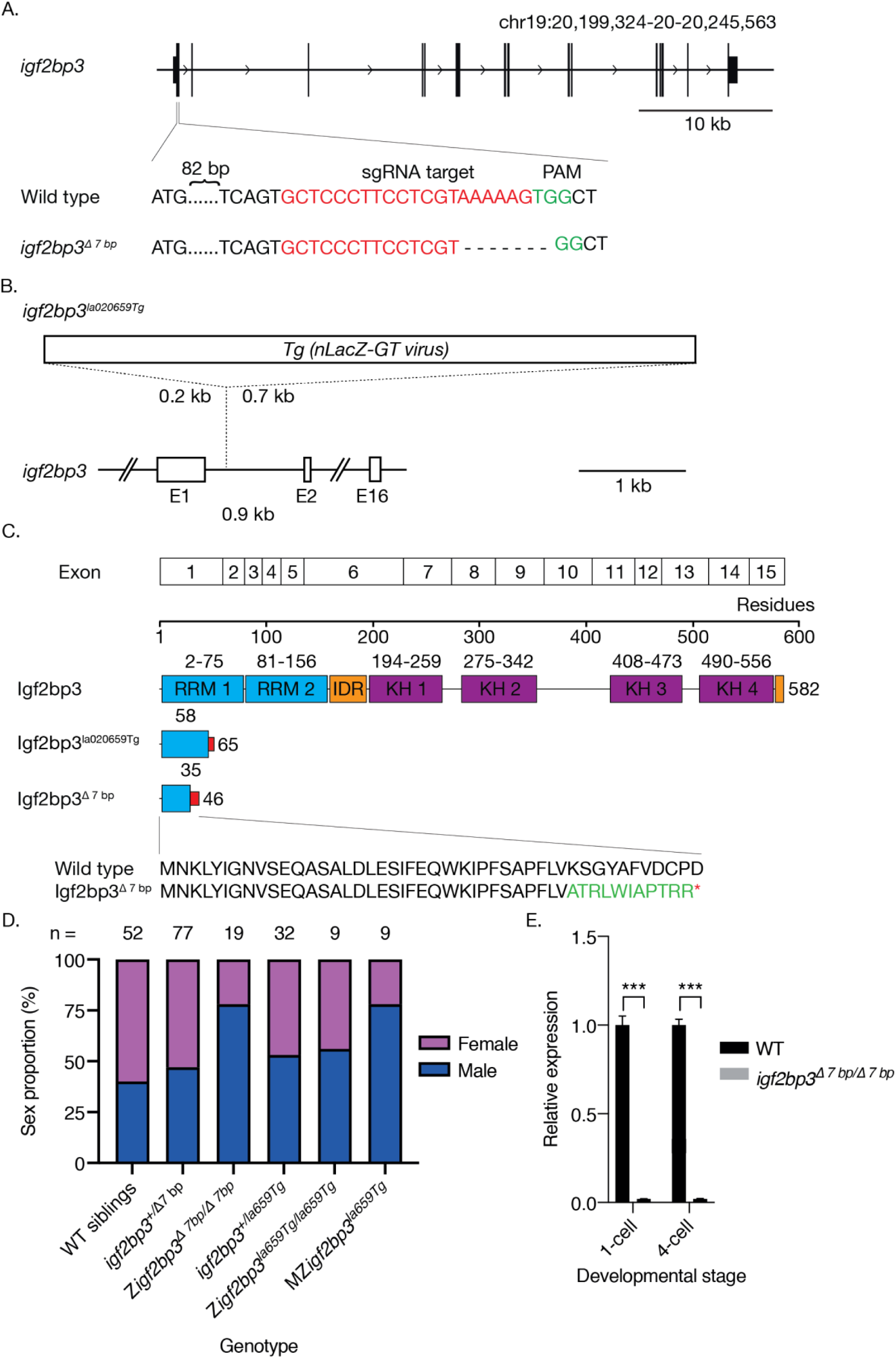
Loss of *igf2bp3* results in skewed male: female sex ratios. A. Generation of the *igf2bp3* mutant allele by Cas9. Schematic showing the igf2bp3 locus on chromosome 19. Exon 1 of *igf2bp3* was targeted by CRISPR-Cas9, resulting in a 7 bp deletion and frameshift. C. The *igf2bp3*_*Δ7*_ mutant is predicted to be protein null. The *igf2bp3*_*Δ7*_ mutation generates a premature stop codon at residue 46 and truncated Igf2bp3 protein. B. Schematic of the *igf2bp3*la020659 Tg with a 6 kb Tg(nLacZ-GTvirus) retroviral insertion in intron 1. D. Loss of igf2bp3 results in skewed sex ratios. Adult zygotic *igf2bp3*_*Δ7*_ and MZ*igf2bp3*la020659Tg mutant fish show a male bias. E. The *igf2bp3*_*Δ7*_ mutation results in reduced *igf2bp3* levels. qRT-PCR at the 1-cell and 4-cell stages showed significant reduction in *igf2bp3* transcript levels.

To generate *igf2bp3* null mutants, we used CRISPR-Cas9 mutagenesis, and identified mutants harbouring a 7 bp deletion within *igf2bp3* exon 1, hereafter referred to as *igf2bp3*_*Δ 7 bp*_ (Fig 1A). The deletion is predicted to result in a frameshift at residue 35 protein that leads to a truncated Igf2bp3 peptide of 47 residues which lacks the RNA-binding domains and the predicted intrinsic disordered regions (Fig 1C), and results in a significant reduction in *igf2bp3* RNA (Fig 1E).

Zygotic *igf2bp3*_*la659Tg*_ grow to adults at Mendelian ratios and appear to be morphological normal. However, progeny of *igf2bp3*_*la659Tg*_ adults (i.e., MZ*igf2bp3* _*la659Tg*_ show biased sex ratios as adults (Fig 1D), with an approximately 3:1 male-female ratio. We also observed a similar male sex bias (3:1) in zygotic *igf2bp3*_*Δ 7 bp*_ deletion mutant adults, suggesting that Igf2bp3 might have important roles in normal germline, gonad and/or sexual development.

### Maternal Igf2bp3 plays a critical role in early development

Although zygotic *igf2bp3* mutants are viable up to adulthood, we observed defects in the progeny of homozygous *igf2bp3* mutant females, i.e., maternal *igf2bp3*_*Δ 7 bp*_ embryos. Gross defects were found across the blastoderm from the 1k-cell stage, resulting in delayed progression through gastrula and lethality in the majority of embryos by 24 hpf (Fig 2A-C).

**Figure 2.**
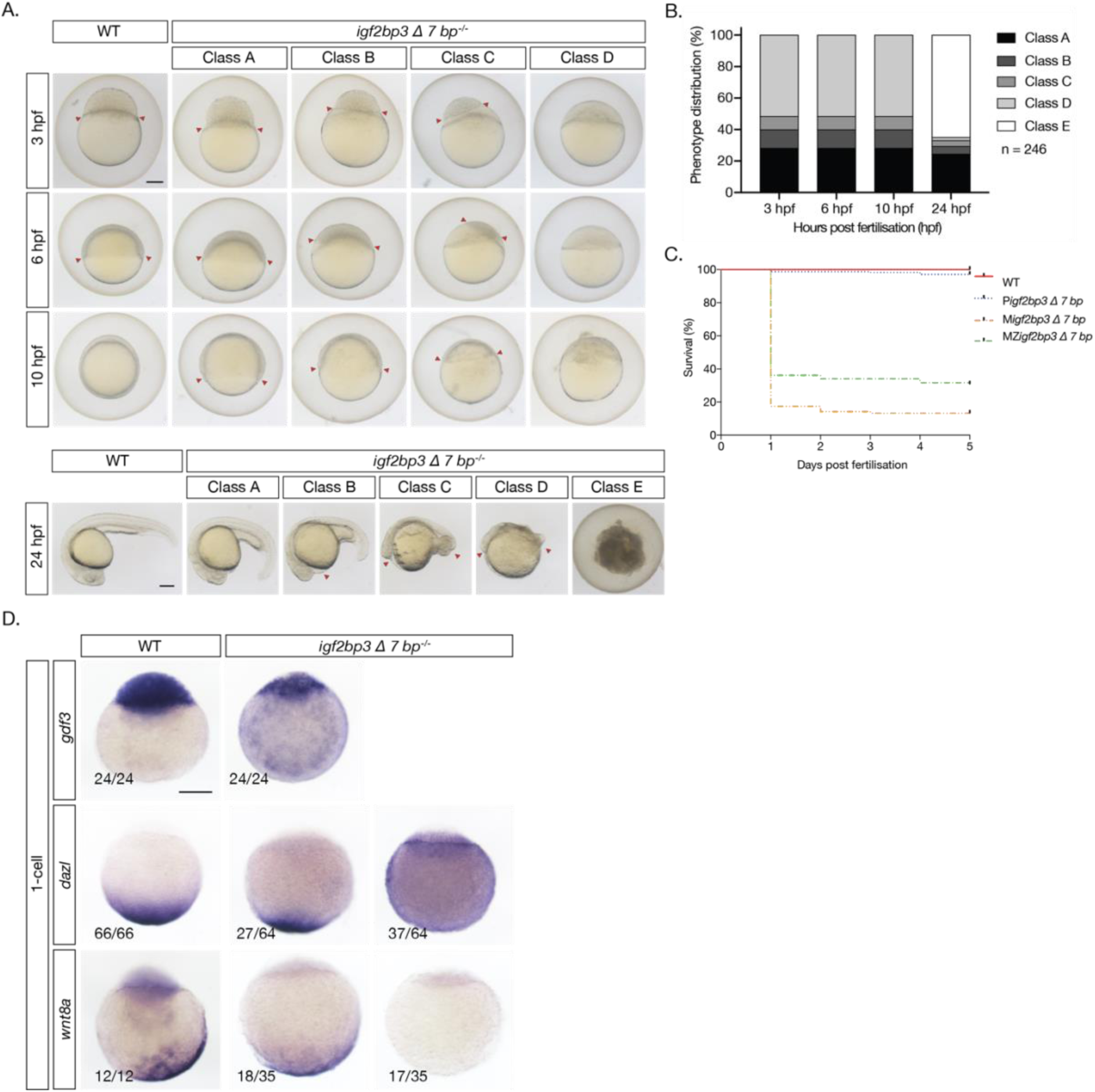
Maternal *igf2bp3* is required for normal embryonic development. **A**. Maternal *igf2bp3* mutants show severe defects by early blastula stages. B,C. The majority of *igf2bp3* mutants do not survive beyond 1 dpf. D. Expression of animal-vegetal markers is reduced or not detectable in *igf2bp3* embryos. Scale bar in A, 200 µm.

As Igf2bp3 is an RNA-binding protein that is involved in the regulation of *vg1* (now *gdf3*) in *Xenopus* and enriched in the Balbiani body, we examined maternal-zygotic *igf2bp3*_*Δ 7 bp*_ embryos at the 1-cell stage for expression of the animal pole marker *gdf3* and vegetal markers, *dazl* and *wnt8a*. MZ*igf2bp3*_*Δ 7 bp*_ embryos showed markedly reduced *gdf3* expression, and either reduced or no *dazl* and *wnt8a* expression (Fig 2D).

### Igf2bp3 is essential for proper germline development

In zebrafish, depletion of primordial germ cells (PGCs) is known to lead to male sex bias To investigate if Igf2bp3 plays a role in PGCs, we examined the germline markers *ddx4, nos1* and *dnd1*, in maternal-zygotic *igf2bp3*_*Δ 7 bp*_ embryos. WISH and qRT-PCRs show that germline markers are reduced in mutant embryos from the 4-cell stage (Fig 3A-B). PGCs are displaced and found in ectopic positions in the blastoderm from the 1k-cell stage and reduced during gastrulation (Fig 3C-D). Ectopic and reduced number of PGCs are observed in the gonadal ridge by 24 hpf (Fig 3E-G) in the maternal *igf2bp3* mutants (progeny from WT males crossed with mutant females, M*igf2bp3*) and MZ*igf2bp3* mutants, but not in the WT and paternal *igf2bp3* mutants (progeny from *igf2bp3* males crossed with WT females, P*igf2bp3*). These defects are also observed in MZ*igf2bp3*_*la361Tg*_ and MZ*igf2bp3*_*la659Tg*_ mutant embryos (Fig Suppl 3).

**Figure 3.**
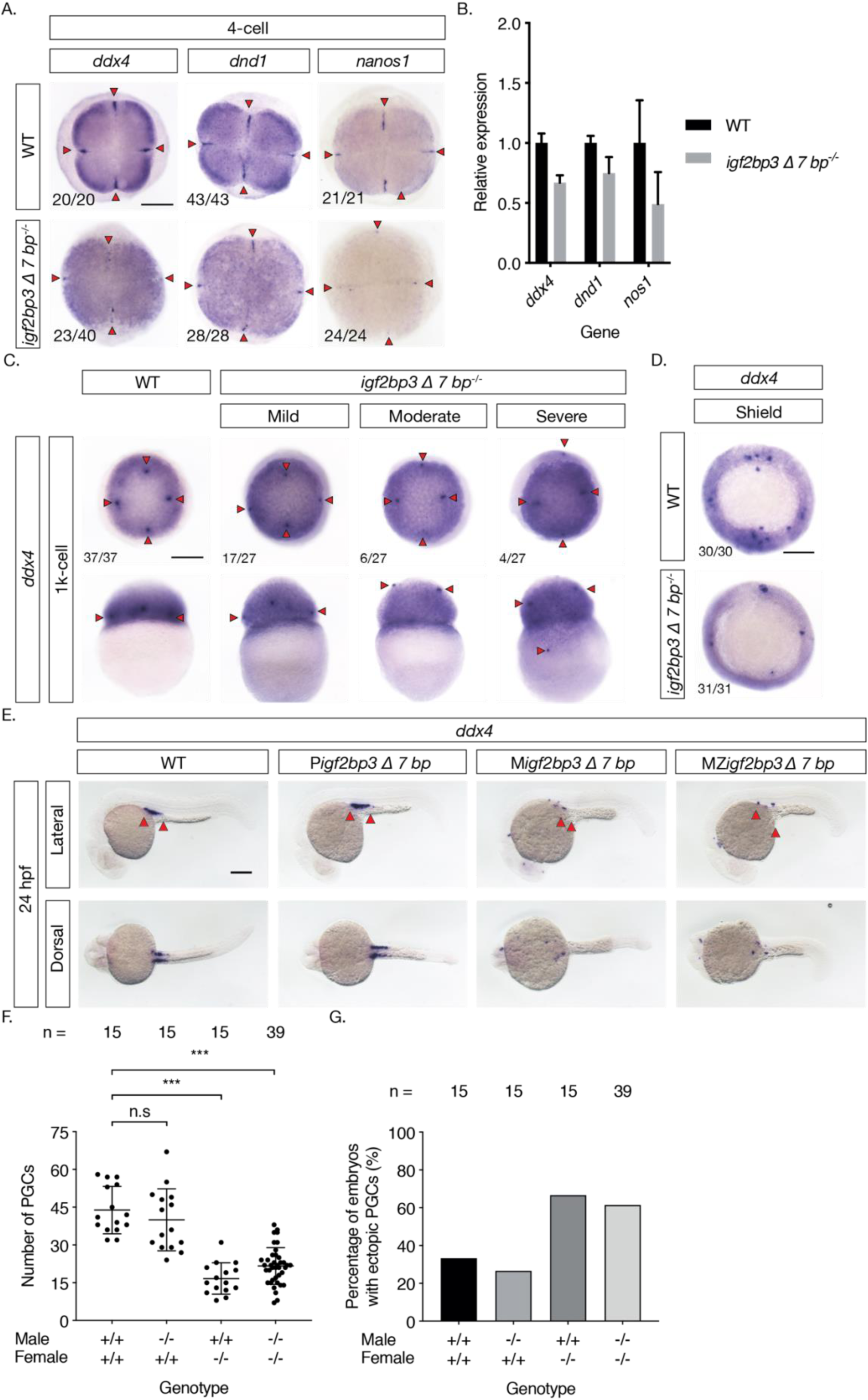
Igf2bp3 is required for normal germ-line development. A. Whole mount in-situ hybridisation in early embryos shows reduced expression of germline markers *ddx4, dnd1*, and *nanos1* in *igf2bp3*_*Δ7*_ mutants embryos. B. qRT-PCR to detect early germplasm markers shows reduced *vasa, dnd1* and *nos1* expression levels in mutant embryos at the 4-cell stage. C. Primordial germ cells in *igf2bp3*_*Δ*_ embryos are ectopically placed at the 1k-cell stage to varying extents ranging from mild or moderate to severe. D. Primordial germ cells are severely reduced or not detected in *igf2bp3*_*Δ7*_ mutants by gastrula stages. E,F. WISH (E) and quantitation (F) of ddx4 positive cells shows reduced and/or ectopic primordial germ cells in 24 hpf maternal *igf2bp3*_*Δ7*_ (M*igf2bp3*_*Δ7*_) and maternal-zygotic *igf2bp3*_*Δ7*_ mutants (MZ*igf2bp3*_*Δ7*_) compared *to* WT siblings and paternal *igf2bp3*_*Δ7*_ (P*igf2bp3*_*Δ7*_) mutants; p * < 0.05, ** < 0.01, *** < 0.001. G. Loss of maternal *igf2bp3* leads to ectopic primordial germ cells. Bra graph shows the number of embryos with ectopic germ cells in WT, P*igf2bp3*_*Δ*_, M*igf2bp3*_*Δ7*,_ and MZ*igf2bp3*_*Δ7*_ mutants. Scale bar in A and C-E, 200 µm.

**Figure 4.**
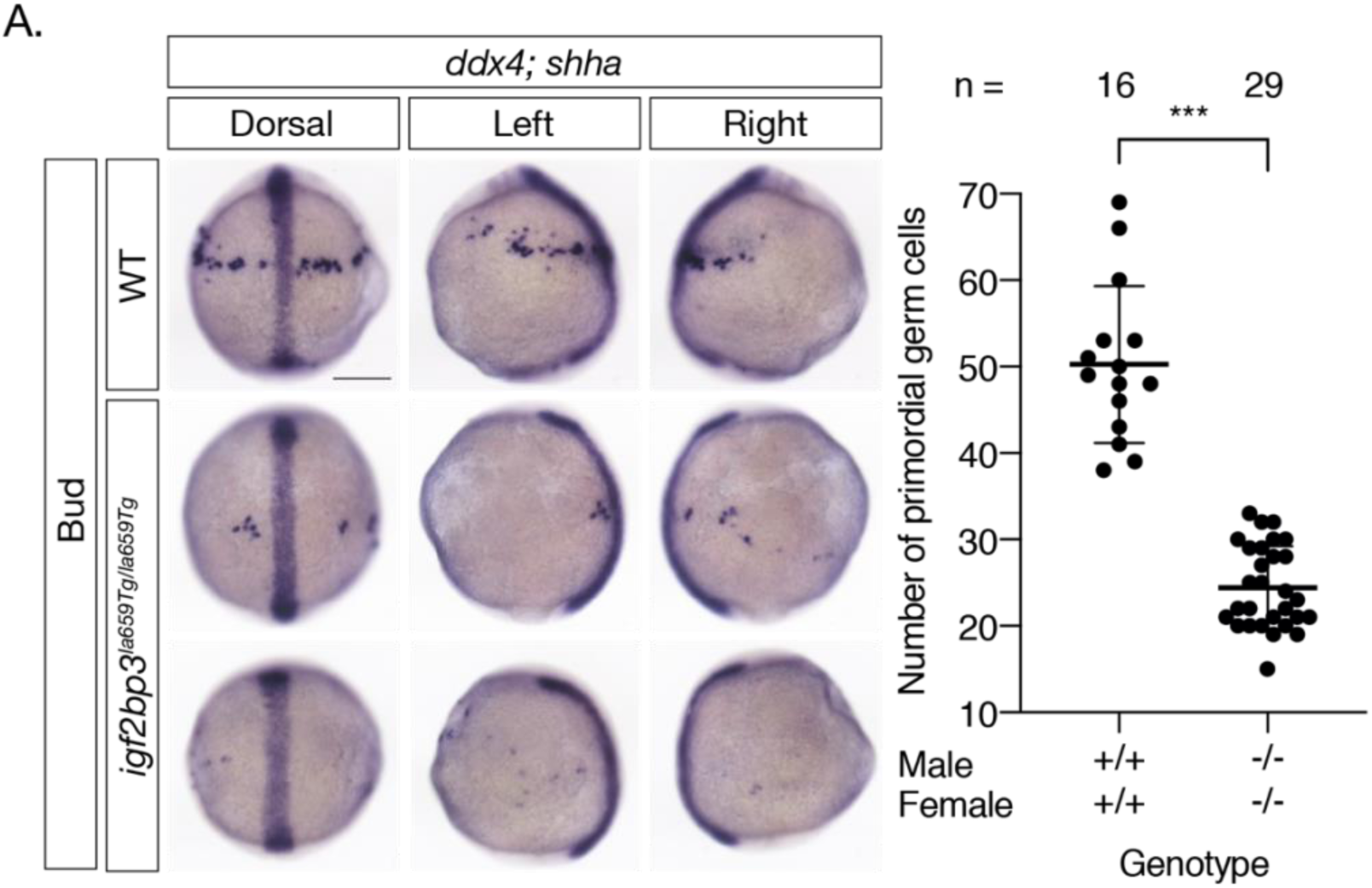
Primordial germ cells are dispersed in the *igf2bp3* mutant. A. PGCs are ectopically located and reduced in M*igf2bp3*_*la020659*_ embryos by Bud stage. PGCs were marked by WISH to detect the germline marker *ddx4*. Scale bar, 200 µm. * p< 0.05, ** < 0.01, *** < 0.001; n=number of embryos analysed.

## Discussion

In this study, we examined the role of the conserved RNA-binding protein, Igf2bp3, in zebrafish, and found that Igf2bp3 is essential for germline development and normal sex ratios in adults, suggesting that *igf2bp3* plays crucial roles in sex determination. We also uncovered a maternal function for Igf2bp3 in early embryonic development.

Previous studies in zebrafish and *Xenopus* found that Igf2bp3 is enriched in the Balbiani body (Bontems *et al*., 2009; Boke *et al*., 2016), a transient structure in oocyte development, where germplasm is initially aggregated and later distributed. The presence of Igf2bp3 as an RNA-binding protein in this structure suggests that it might regulate various RNAs. Consistently, we find that several germline markers, including *ddx4, nanos1* and *dnd1*, are reduced from the 4-cell stage. These findings suggest that Igf2bp3 may act as a stability factor for germplasm RNAs.

Furthermore, various germplasm RNAs are critical for survival, proliferation or migration of the PGCs to ensure they remain distinct from the soma. For example, *dnd1* RNA acts as a survival and migration factor for the zebrafish PGCs (Weidinger *et al*., 2003). The reduction of germline RNAs correlates with the aberrant germline phenotype we have observed, such as mislocalisation and depletion of PGCs, as well as abnormal migratory behaviour when live PGCs are tracked.

A recent study examining the zebrafish transcriptome in *igf2bp3* mutants reports that *igf2bp3* acts as a stability factor for maternal RNAs in development (Ren *et al*., 2020), However, this study has focused on the misregulation of a subset of target mRNAs in their *igf2bp3* mutants. By contrast, we have observed additional phenotypes that were not previously been reported, such as a male sex bias in zygotic mutants. Therefore, *igf2bp3* might have crucial roles in further processes that are not strictly maternal.

The mechanism that Igf2bp3 acts through to regulate the germline remains unknown. However, many proteins associated with m6A modifications, known as m6A readers, have been shown to be important for fertility and germline development, such as Ythdf2 (Zhao *et al*., 2017), Ythdc2 (Hsu *et al*., 2017; Soh *et al*., 2017; Jain *et al*., 2018) and a closely related *Drosophila* homolog, Bcgn (Gateff, 1982). Igf2bp proteins were found to be associated with m6A modifications (Huang *et al*., 2018). the phenotype we have observed in the Igf2bp3 mutants is concordant with the literature and implicates Igf2bp3 to be part of a conserved network of proteins involved in RNA regulation with potentially conserved functions.

---

**Figure Supplement 1.**
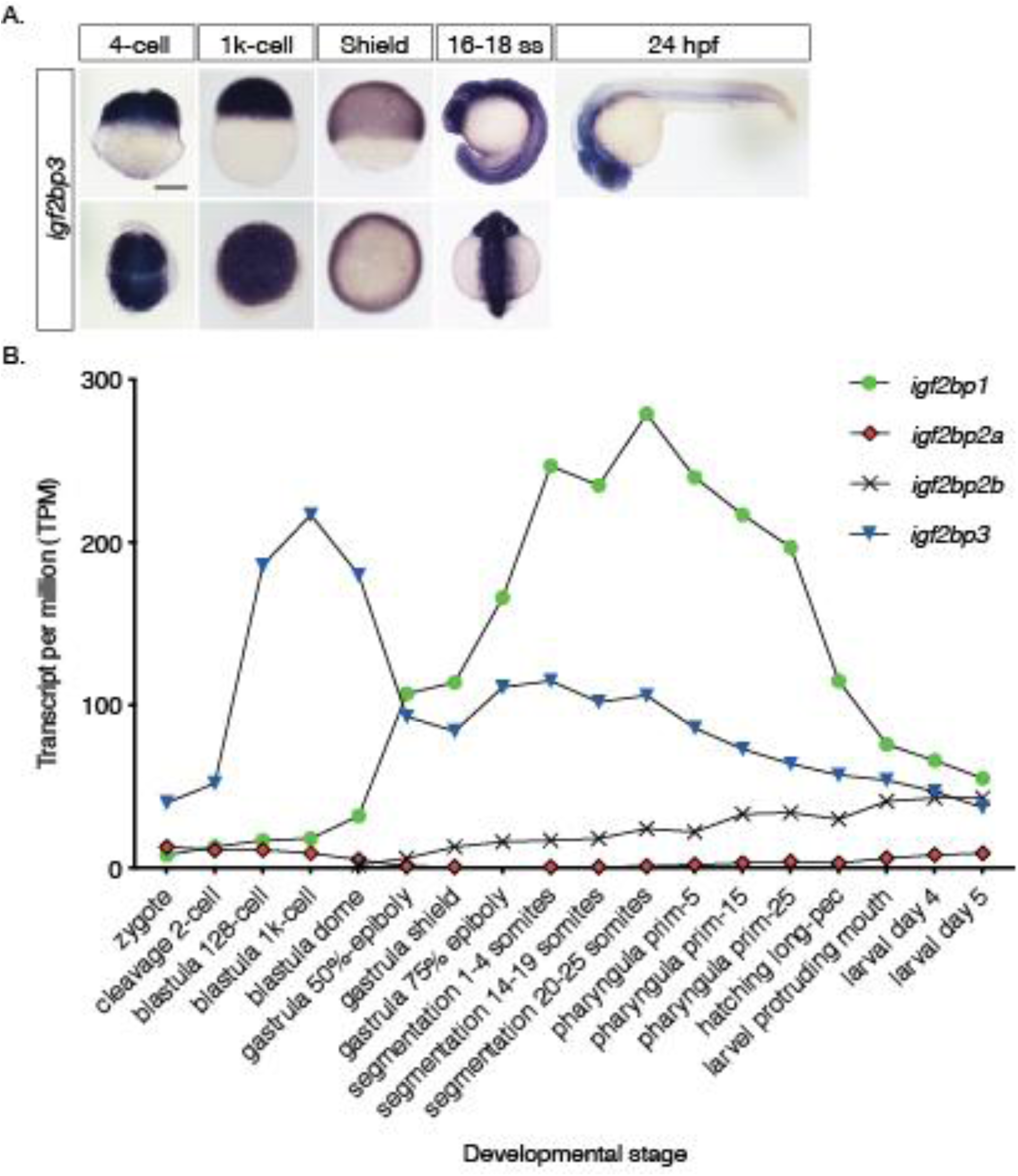
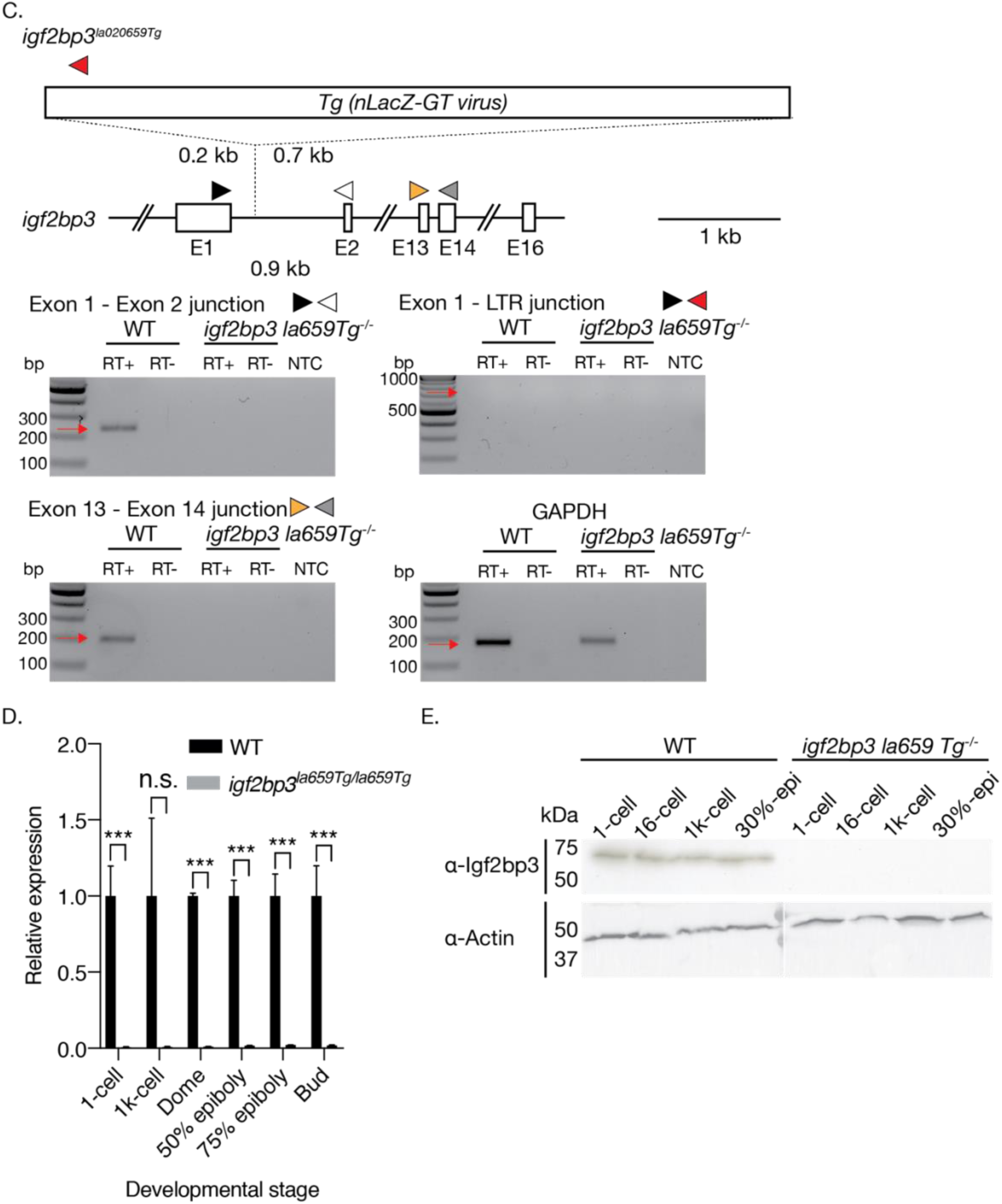
*igf2bp3* is expressed ubiquitously during early development. A. Whole *in-situ* hybridisation to detect *igf2bp3* in zebrafish embryos at cleavage, blastula, gastrula, somitogenesis and 24 hpf. B. RNA-seq expression data for all four zebrafish *igf2bp* genes during zebrafish development shows that *igf2bp3* is expressed throughout zebrafish development. Data from (White *et al*., 2017). C, D. MZ*igf2bp3*la020659Tg mutant embryos show significantly reduced or undetectable expression of *igf2bp3*. RT-PCR (C) using exon spanning primers for *igf2bp3* and qRT-PCR (D) at performed at six early developmental stages, show reduced products in mutant embryos relative to *gapdh*. E. Igf2bp3 protein is not detectable in MZ*igf2bp3*la020659Tg embryos. Western blots on cleavage to early gastrula embryos show a 60 kD band in wild type (WT) lysates. No bands for Igf2bp3 are detected in MZ*igf2bp3*la020659Tg lysates. Scale bar in A, 200 µm.

**Figure Supplement 3.**
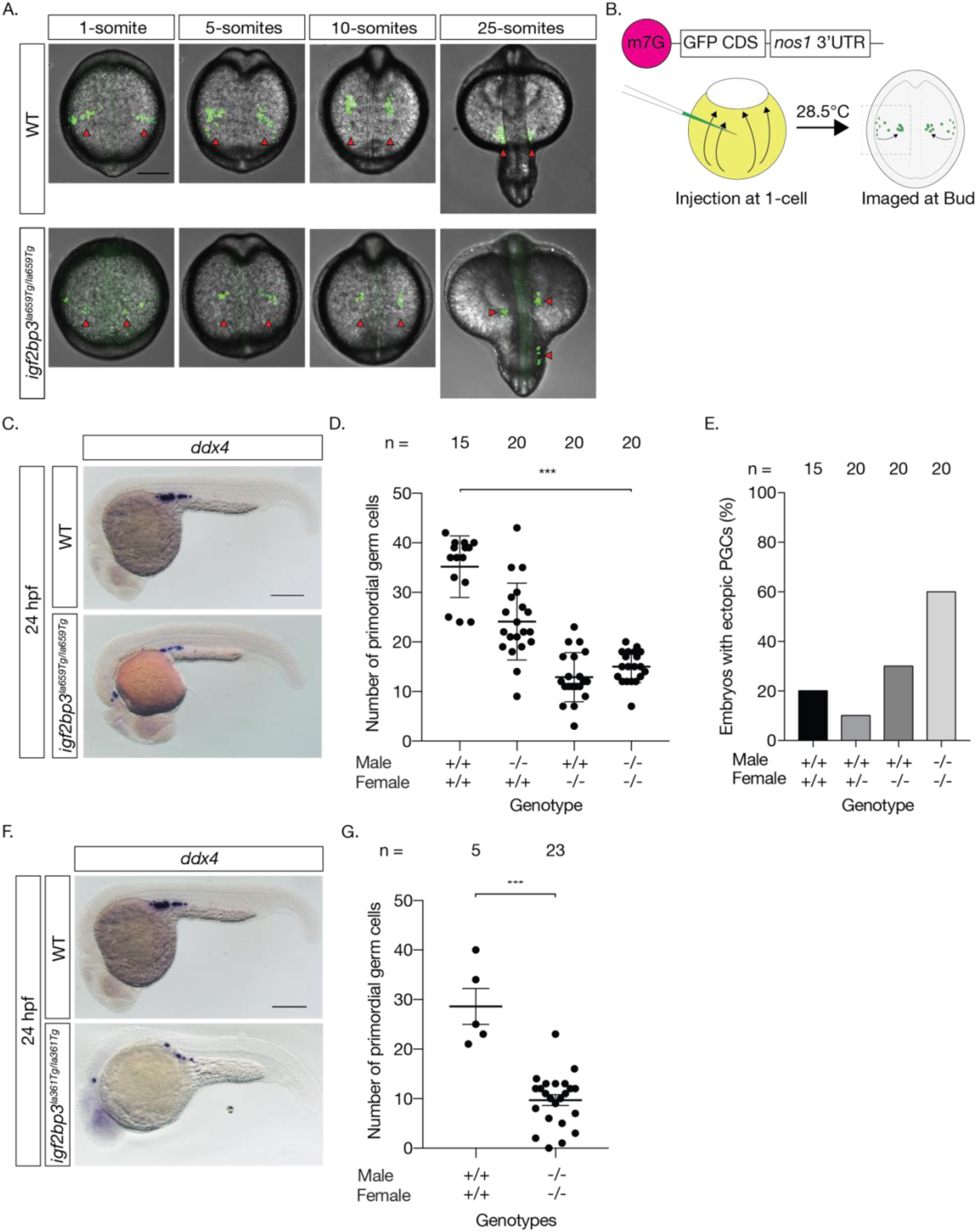
The germline is misregulated in *igf2bp3* mutants. A. Live imaging of PGCs in embryos injected with GFP-nos3’UTR reporter mRNA shows reduced PGC numbers in *igf2bp3la659Tg/la659Tg* mutants compared to WT embryos at 1-somite, 5-somite, 10-somites and 25-somites. C-E. The germline is mis-regulated in *igf2bp3la659Tg/la659Tg* mutant embryos with PGC numbers reduced and ectopically located in the MZ *igf2bp3la659Tg/la659Tg* mutants (C, E), and reduced in the M*igf2bp3la659Tg/la659Tg* and MZ *igf2bp3la659Tg/la659Tg* (D). F. The germline is also mis-regulated in a second transgenic insertion mutant line, *igf2bp3 la361Tg* with reduced PGCs in MZ *igf2bp la361Tg* mutants (F, G). Scale bars in C and F, 200 µm; * p < 0.05, ** < 0.01, *** < 0.001.

**Figure Supplementary 4.**
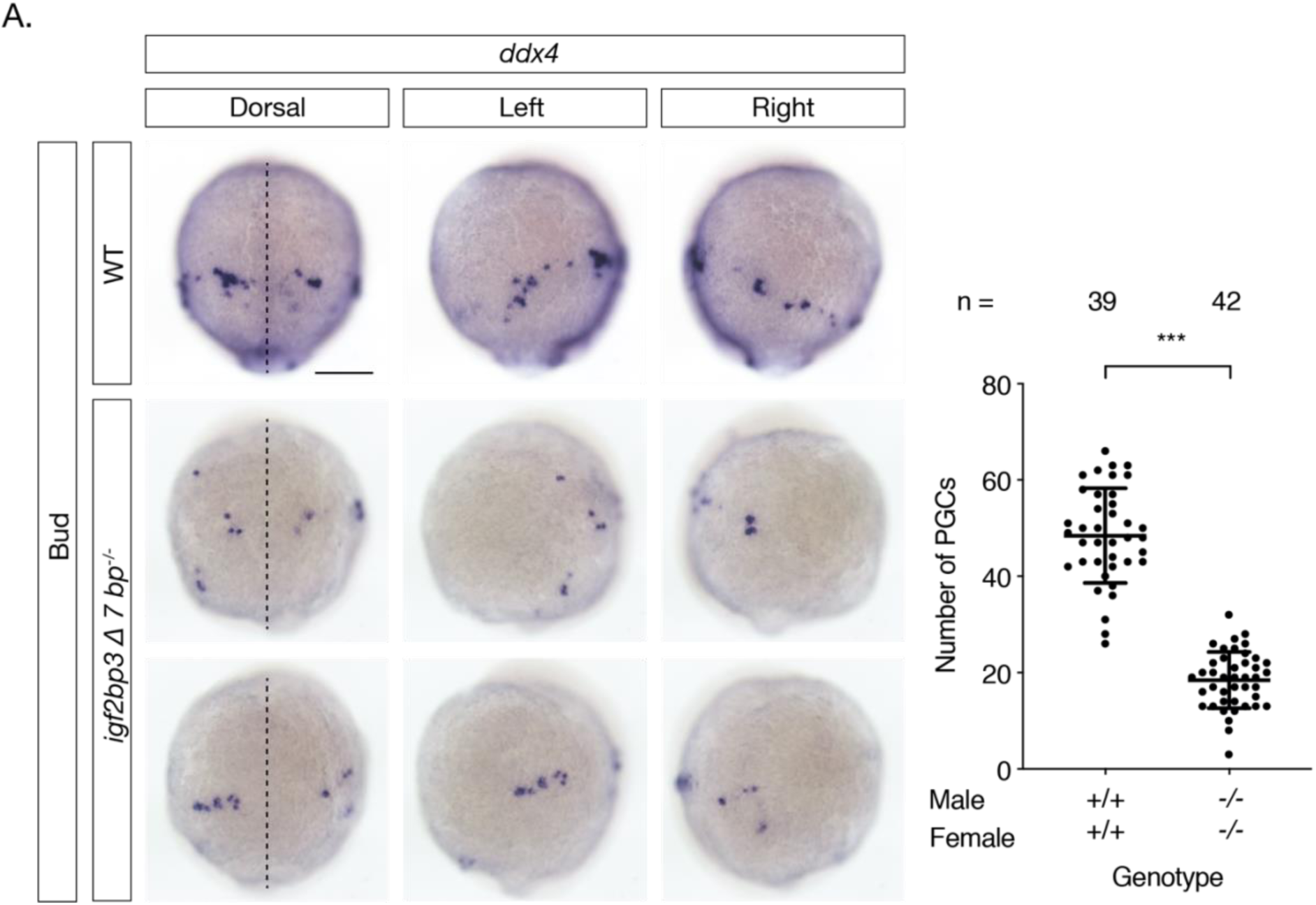
PGC migration and numbers are reduced in *igf2bp3*_*Δ7*_ mutants. AWISH to detect ddx4 shows that PGCs are ectopically located and reduced in the *igf2bp3*_*Δ7*_ embryos by Bud stage. Dashed line indicates midline; Scale bar, 200 µm. * p< 0.05, ** < 0.01, *** < 0.001.

## Supplementary Methods

qRT-PCR primers

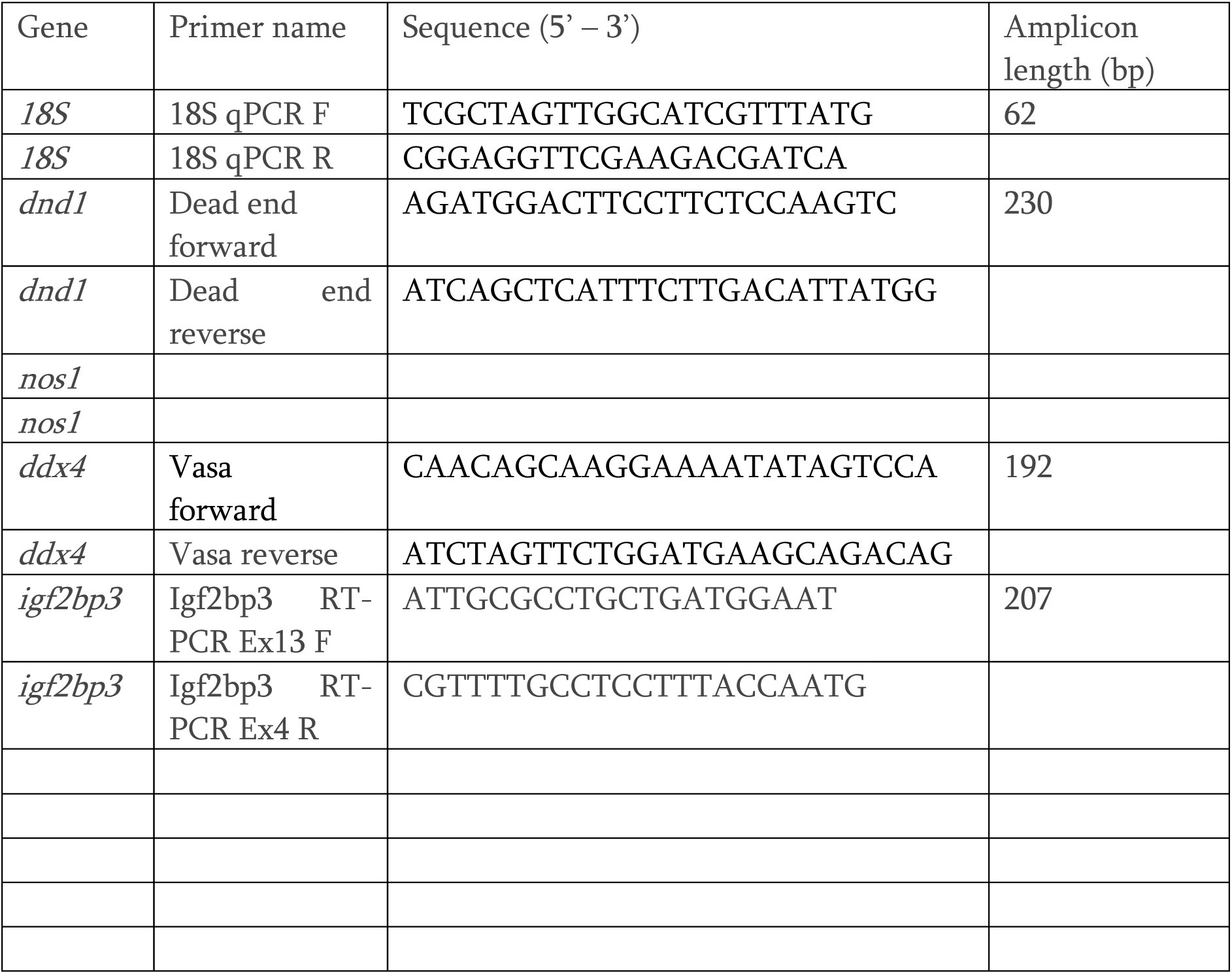

Genotyping primers

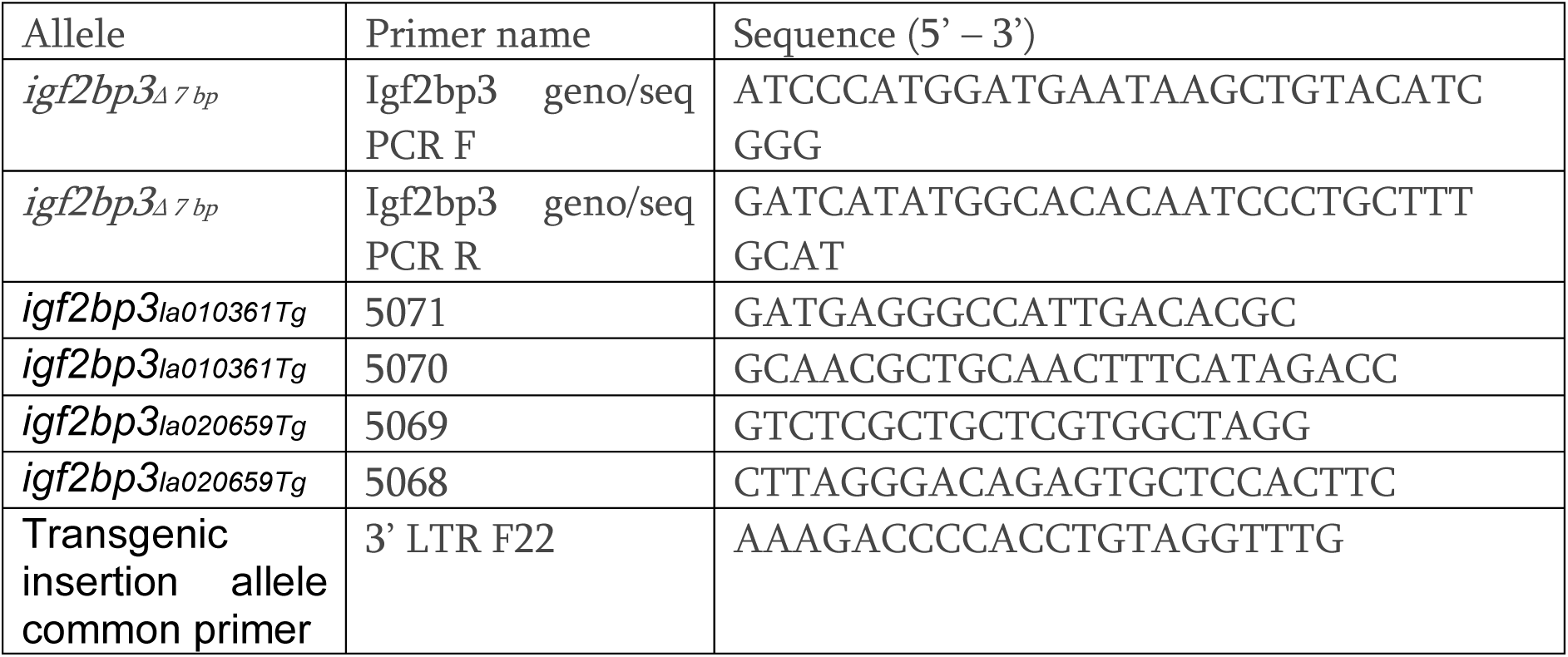

## Aptamers for Tobramycin pull down

### Aptamer sequence

ATGCTAGCGGGAGAAGACGACCGACCAGAATCATGCAAGTGCGTAAGATAGTCGCGGGCCGGGAAAAAAAAAAGGCTTAGTATAGCGAGGTTTAGCTACACTCGTGCTGAGCCAAAAAAAAAAGACCGACCAGAATCATGCAAGTGCGTAAGATAGTCGCGGGCCGGGTATGTGCGTCTG

### ndr1/sqt 5’UTR: aptamer: ndr1/sqt 3’UTR

ACGAGCTTTATTTCAATAACTGCGTGTGGATTATTACCTTGATTTGACATGTTTTCCTGCGGGCTCCTGAGCGTAGTTTTGGCCCTTATGCTAGCGGGAGAAGACGACCGACCAGAATCATGCAAGTGCGTAAGATAGTCGCGGGCCGGGAAAAAAAAAAGGCTTAGTATAGCGAGGTTTAGCTACACTCGTGCTGAGCCAAAAAAAAAAGACCGACCAGAATCATGCAAGTGCGTAAGATAGTCGCGGGCCGGGTATGTGCGTCTGGATCCTATTGGCAAGATGGTCATGAGACACCATGAAGGCATGGTTGTTGCAGAATGCGGCTGCCACTGATTCTTCAAACCCCAAAGGAACTCAACTCTAGCACTTTGGATATGCTCCTTGACCCCAAAAATATGTATTTAAGAAAAACTGCTGTCAATTATTCCCACTTGAAATTATTATGGTTTCCTGCACTGAGGCACCTGGATAACTTGATGCTATTATTGAAAGCTTTGCGTGTTTGCCTTATCTGTAAATAGTAGAGTATGTAAATTACCAAATGTAATAAAATGTTTTCATAATGTTTAAAAAAAAAAAAAAAAAA

### Aptamer: ndr1/sqt 3’UTR

TATGCTAGCGGGAGAAGACGACCGACCAGAATCATGCAAGTGCGTAAGATAGTCGCGGGCCGGGAAAAAAAAAAGGCTTAGTATAGCGAGGTTTAGCTACACTCGTGCTGAGCCAAAAAAAAAAGACCGACCAGAATCATGCAAGTGCGTAAGATAGTCGCGGGCCGGGTATGTGCGTCTGGATCCTATTGGCAAGATGGTCATGAGACACCATGAAGGCATGGTTGTTGCAGAATGCGGCTGCCACTGATTCTTCAAACCCCAAAGGAACTCAACTCTAGCACTTTGGATATGCTCCTTGACCCCAAAAATATGTATTTAAGAAAAACTGCTGTCAATTATTCCCACTTGAAATTATTATGGTTTCCTGCACTGAGGCACCTGGATAACTTGATGCTATTATTGAAAGCTTTGCGTGTTTGCCTTATCTGTAAATAGTAGAGTATGTAAATTACCAAATGTAATAAAATGTTTTCATAATGTTTAAAAA AAAAAAAAAAAAA

